# Matrix-seq: An adjustable-resolution spatial transcriptomics via microfluidic matrix-based barcoding

**DOI:** 10.1101/2022.08.05.502952

**Authors:** Haifeng Zhao, Geng Tian, Aihua Hu

**Affiliations:** Intelligent Healthcare Co., Ltd., Beijing, 100023, China; Geneis Beijing Co., Ltd., Beijing, 100102, China; Beijing Children’s Hospital, National Center for Children’s Health, Capital Medical University, China

**Author notes:** Corresponding author: Haifeng Zhao.

## Abstract

Spatial transcriptomics technology complements the spatial information lost in single-cell RNA sequencing, which enables visualization and quantitative analysis of transcriptomics of cells in tissue sections. Although this technology is a promising tool to study complex biological processes, its popularization is limited by cumbersome barcoding steps. We presented a microfluidics-based barcoding strategy called Matrix-seq, which gets rid of both precision instruments and the *in situ* indexing. The deterministic barcoding matrix is fabricated by the crossflow of Barcode-X and Barcode-Y. The overlapping areas (spot) formed deterministic barcoding primers (Barcode-X-Y) via the ligation reaction. Matrices with different spot size (ranging from 10 to 50 μm), which was decided by the width of microchannels, were fabricated and then applied to a mouse main olfactory bulb section and a mouse brain section. While maintaining high performance and resolution, this technology greatly reduces the technical threshold and cost of spatial barcoding. As a result, Matrix-seq can be rapidly applied in various fields including developmental biology, neuroscience and clinical pathology.

**Teaser:** Matrix-seq provides an orthogonal microchannel-based barcoding strategy for adjustable-resolution spatial transcriptomics.

## Introduction

Single-cell RNA sequencing (scRNA-seq), which provides unbiased high-throughput gene expression information and reveals the unexpectedly large degree of heterogeneity in cell types and cell states in bulk tissue, has become a key technology in biomedical research and disease diagnosis in the past decade(*1-5*). The disassociation process during the sample preparation inevitably leads to the loss of spatial information of individual cells, which severely limits the understanding of tissue complexity in normal physiology or under perturbation. Specifically, scRNA-seq is weak in unveiling cell status, in which cell location plays important roles, such as cell-cell interactions, tissue microenvironments, and so on. Spatial transcriptomics technology complements the spatial information lost in tissue dissociation, which enables visualization of transcriptomics in tissue sections(*6*). Several remarkable methodologies have been developed, including image-based and next generation sequencing (NGS)-based technologies(*7*). Image-based methods relies on multiplexing fluorescent probes, including in situ hybridization (ISH) or sequencing (ISS)(*8*). Although it is possible to detect gene expression in the whole transcriptome scale recently(*9*), this technology still has a high technical threshold, relying on high-sensitivity single-molecule fluorescence imaging systems and a lengthy, repeated and sophisticated workflow(*8, 10*). The methods powered by NGS provide unbiased, genome-level, high throughput and cost-efficient analytical solutions, and the difference among NGS-based methods is mainly in the spatial barcoding strategies. In the early stage, laser capture microdissection (LCM) was used to segment different regions to preserve spatial information before dissociation(*11, 12*). Although LCM is a robust technology with cell-level cutting accuracy, it is very labor intensive, which limits the throughput of samples. In recent years, various technologies for spatial barcoding have emerged. In terms of barcoding strategies, they can be divided into deterministic and random barcoding. The former marks deterministic barcoding primers at a fixed position, while the latter randomly synthesizes the barcoding primer and then performs decoding and indexing by in-situ sequencing. The typical techniques are spatial transcriptomics (ST) and Slide-seq(*6, 13*). In terms of ligation between RNA and barcoding primers, it can be divided into capture- and labeling-based strategies. The former releases the RNA in the tissue section and captures them by the primers on the solid-phase substrate (slide, silicon wafer or beads), and the latter directly labels the primers into the tissue. Typical techniques are HDST and DBiT(*10, 14*). Although Visium (10x Genomics) was the first to be commercialized, the high cost, low yield and low resolution of the barcoding strategy limited its wide-scale promotion.

It is highly desirable to develop a barcoding strategy with high-spatial resolution, high sensitivity, and low cost. In addition, it should be easy to store and has no requirement for special instruments, to facilitate the usage of researchers without special training. Due to an imbalance between resolution, efficient area and cost, ideal protocols can freely adjust the resolution and area to match cell density and size in different samples. Essentially, NGS-based techniques are two-dimensional encoding and indexing of transcripts in tissue sections. Similar addressing problems exist widely in chess sports, mathematics and computer sciences. Row-column addressing is one of the most classic schemes. As early as two thousand years ago, the ancients in China used row-column to describe the position of chess pieces on the chessboard. Prof. Fan’s group took the lead in applying this idea to spatial omics (DBiT-seq)(*10*). They used row-column microfluidics for delivering barcoded probes directly into tissues and can simultaneously label transcripts and selected protein targets. Compared with the independent and serial barcoding of each spot, the barcoding based on row-column microfluidics can be carried out in parallel, simplifying thousands of dispensing processes into two injections, which greatly reduces the labor and requirements for the capacity of the barcode library. Although in-situ labeling achieves high sensitivity and compatibility with multi-omics(*15*), the risk of clogging and leakage due to the soft nature of the tissue section significantly reduces the yield and causes the inhomogeneity between microchannels. These defects often require sequencing analysis to detect, which is clearly unacceptable for the analysis of precious clinical samples and poses a challenge to quality control.

Here, we developed a different barcoding strategy called the matrix-based spatial transcriptomics (Matrix-seq), which applies row-column microfluidics to fix and ligate deterministic barcoding primers to slides. The primers are produced by two sets of parallel microfluidic channels (10, 25, or 50 μm in width), defined as Microchannel-X and Microchannel-Y, which are orthogonal to each other. The Barcode-X (BC-X, *X*_1_ to *X*_70_) transported in the Microchannel-X was modified in the functionalized slide, and then the crossflow of Barcode-Y (BC-Y, *Y*_1_ to *Y*_70_) in the Microchannel-Y yielded a matrix of deterministic barcoding primers (*X*_1_*Y*_1_ to *X*_70_*Y*_70_) on the overlapping area via the ligation reaction. The resolution of matrix (width of spots and gaps) was determined by the width of the microchannels and walls, which brings a high degree of flexibility in the design of its resolution, coverage and field of view. The fabricated slides can be transported and cryopreserved in the same way as conventional DNA chips. When in use, the users paste the tissue section on the barcoded area, and then performs fixation, H&E staining, permeabilization, digestion and reverse transcription (RT) in sequence. The barcoded cDNAs were collected into a tube. After PCR amplification and tagmentation, the library was prepared for NGS sequencing. We demonstrated a matrix arrayed by 70 × 70 spots with 50/30 μm width of microchannel/gap for spatial transcriptomic mapping of main olfactory bulb (MOB) and half brain of mouse. Matrix-seq faithfully identified the anatomical structures and detected the whole transcriptome. While retaining the advantages (efficiency, high-resolution, simple and easy-operation) of row-column barcoding, Matrix-seq divides the preparation process into the slide part and tissue part, which makes the quality control of barcoded matrix very convenient and allows researchers without special training to fabricate these slides boldly without worrying about the loss of precious samples. This technology provides a full-process solution from barcoding to analysis, lowering the threshold of spatial transcriptomics to an unprecedented level, which can be adopted in any laboratory with a pipette, cryostat, microscope and PCR amplifier.

## Results

### Workflow of Matrix-seq

The workflow of Matrix-seq was described in Fig. 1, which can be divided into three parts, i) preparation of Matrix-slide, ii) spatial barcoding of tissue sections and iii) data analysis. Microchannel-X containing 70 parallel microchannels was placed on the functionalized glass slide. The structure of this assembly was shown in Fig. 2A, where Microchannel-X was attached to a standard glass slide (75 mm × 25 mm). The set of BC-X solutions were introduced in the inlet wells and driven in parallel by a vacuum pump, which can be accomplished in several minutes. The BC-X is composed of an amino in 5’ end for binding to the epoxy-modified slide, a PCR handle, a deterministic BC − *X*_i_ (i=1-70), and a ligation linker. The binding reaction between the BC-X and slide was conducted in the microchannel under the condition provided by the manufacturer. Then, the Microchannel-X was replaced by Microchannel-Y. The microchannels in Microchannel-Y are perpendicular to those in Microchannel-X. The BC-Y contains a ligation linker, a deterministic BC − *Y*_j_ (j=1-70), a unique molecular identifier (UMI), and a series of T (poly T) for mRNA capture. The regents mixed by BC-Y solution, T4 ligase and a complementary ligation linker were driven into Microchannel-Y for in situ ligation at the overlapping area, resulting in a matrix consisted of 4900 square spots. Each spot was indexed by the spatial coordinate (row-column, *X*_i_ − *Y*_j_) and contains a deterministic barcoded primer (BC − *X*_i_*Y*_j_). Prepared slides can be easily stored and transported until used. When barcoding the tissue, a frozen tissue section (10 μm) was loaded onto the Matrix-slide, followed by fixation, H&E staining, and permeabilization to release RNAs, which was captured by barcoded primers. After RT and amplification, amplified and barcoded cDNA were collected and used as the template for library preparation and NGS. Based on the strict correspondence between spatial coordinates and barcoding sequences, transcript data can be easily classified into the corresponding spatial position. This technique builds a bridge between classical bioinformatics analysis of transcripts and pathologic analysis of H&E images to reveal the gene expression in individual spots and shows the corresponding tissue morphology synchronously. The key reagents, sequences of barcodes and design of microchannels were summarized in Tables S1 and S2 and shown in fig. S1, respectively.

**Fig. 1.**
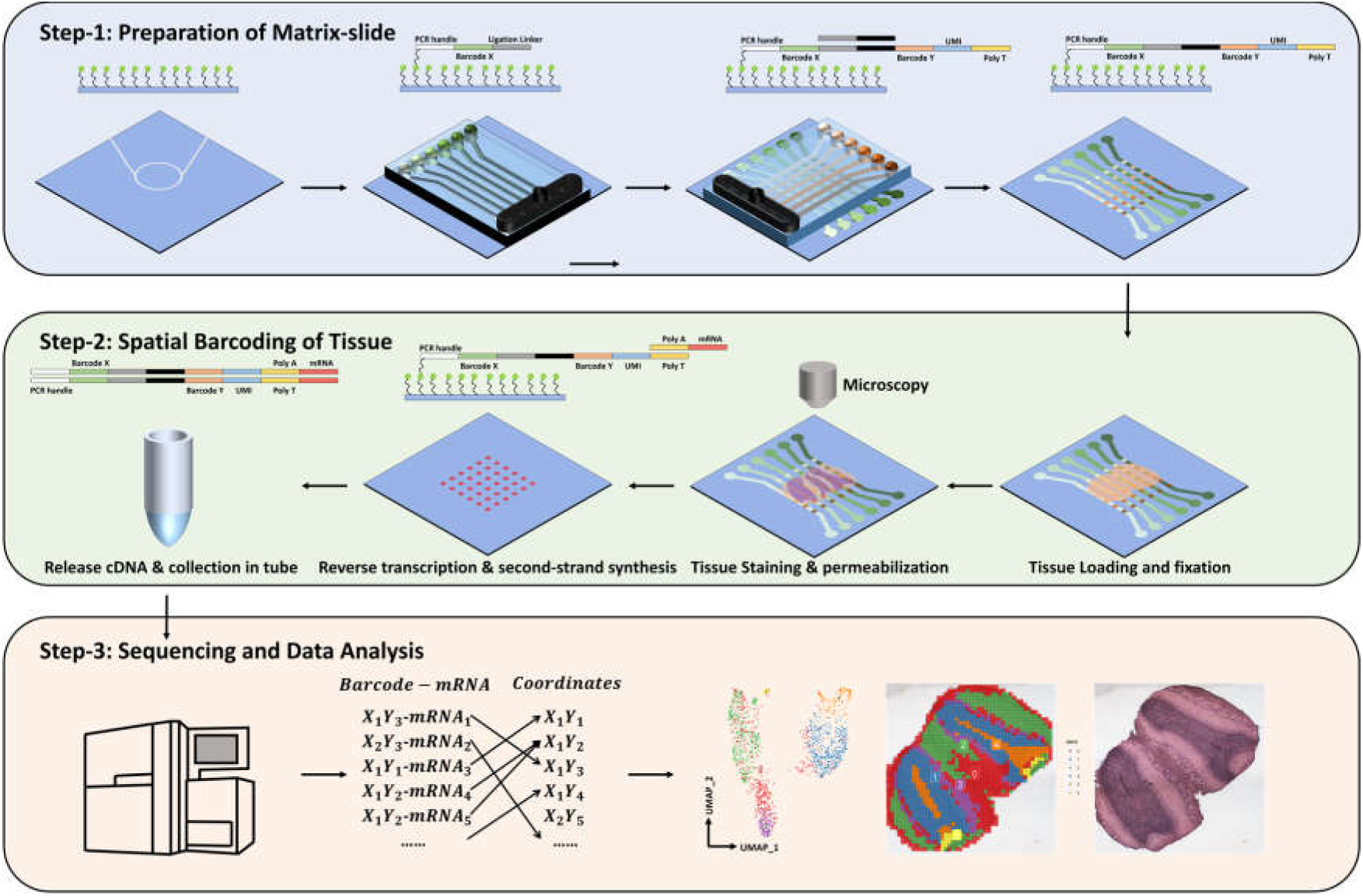
Matrix-seq pipeline. Step-1, preparation of the Matrix-slide. The barcoded spots are fabricated by two set of parallel microchannels with binding and ligation reaction. Step-2, spatial barcoding of tissue. After fixation and H&E staining, The RNAs released from the tissue section by permeabilization are in situ capture by the barcoded primers. Followed by RT and second-strand synthesis, the released cDNA is collected in tube for library construction. Step-3, sequencing and data analysis. a spatial mRNA map is reconstructed by matching the spatial barcodes to spatial spots. The transcript map can be correlated to the H&E image to identify the fine structures.

**Fig. 2.**
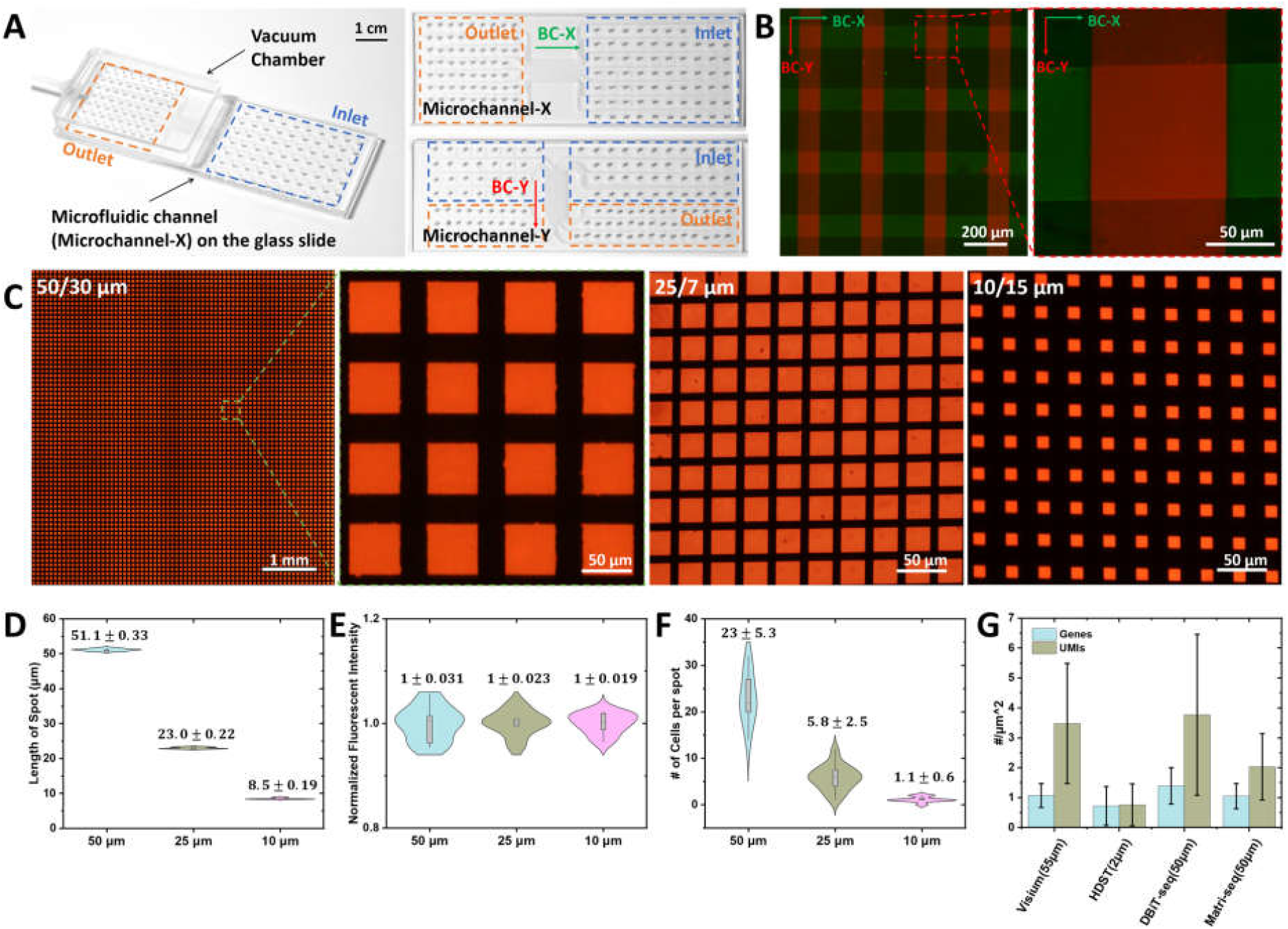
Evaluation of barcoding process and performance. **(A)** Structure of Matrix barcoding platform. The platform is consisted of a vacuum chamber, microfluidic chip and functionalized glass slide. The images of Microchannel-X and Microchannel-Y are shown in left. The regions of inlets and outlets are marked by blue and orange dashed rectangles. The direction of BC-X and BC-Y are pointed by green and red arrows. **(B)** The merge fluorescent image demonstrates the distribution of BC-X (green) and BC-Y (red). The detail image shows the overlapping area exhibits stronger fluorescence and sharper boundaries. **(C)** The fluorescent images of matrices consisted of spots of 50, 25 and 10 μm. The gaps between the spots are 30, 7 and 15 μm, respectively. **(D-F)** Qualification of the length of spots, uniformity of fluorescence and the number of cells per spot. **(G)** Evaluation of capture performance. Normalized gene and UMI count of Matrix-seq was compared to Visium, HDST and DBiT. The fresh frozen tissues of MOB were used in Matrix-seq, HDST and Visium. The fixed embryonic brain was used in DBiT.

### Evaluation of microfluidic matrix-based barcoding

To benchmark the barcoding process, fluorescent-labeled BC-X (green, fluorophore fluorescein isothiocyanate (FITC)) and BC-Y (red, Cy3) were used to visualize the spatial distribution of oligonucleotides. The fluorescent matrix consisted of row and column fluorescent bands was generated as expected (Fig. 2B). The overlapping areas exhibited strong red fluorescence indicating the efficiency of ligation reaction. Ideally, the red fluorescent of BC-Y should not be present on non-overlapping areas. We suspect this is related to non-specific adsorption. The clear boundaries indicate that this method exhibits high precision and robustness, which is difficult to achieve on the tissue surface due to potential diffusion effect and its unevenness and softness. Further, we fabricated matrices with different spot size (50, 25 and 10 μm), which was visualized by fluorescence-labeled probes (red, Cy3) via RT reaction to evaluate the spatial accuracy of this method (Fig. 2C). Among them, the matrix consisted by 70 × 70 spots of 50 μm was prepared for subsequent tissue experiments, while the spots of 25 μm and 10 μm were generated respectively to demonstrate the superiority of this method in coverage and resolution. Coverage is defined here as the ratio of the effective area (spots) to the total matrix area, which is determined by the size of spots and gaps. Higher coverage can capture more released transcripts and make the map mote complete, but it requires higher spatial accuracy of lattice generation to prevent the confusion of barcoding. For example, the ST technology used circular spots with a diameter of 100 μm and a center-to-center distance of 200 μm and Stereo-seq used a grid-patterned array of spots with approximately 220 nm in diameter and a center-to-center distance of 500 nm, which means their coverage is about 19.6% and 15.2%(*6, 16*). Even after optimization for commercialization, the coverage of Visium with a diameter of 55 μm and a center-to-center distance of 100 μm is still about 15.9%(*17*). Compared to them, Matrix-seq has solid walls to separate different barcode solution, which brings excellent stability, robustness and accuracy. As a demonstration, we produced a matrix consisted of square spots of a diameter of 25 μm and a center-to-center distance of 32 μm with coverage up to 61%. Due to the high precision of the photolithography process, high-resolution spots of 10 μm can also be produced by this method with high yield and uniformity. Based on fluorescent images of RT, we further characterized the uniformity of spot size and fluorescent intensity, which represent the processing accuracy and uniformity of primers, as shown in Fig. 2D and E. The point where the fluorescence value drops to 10% of that of spot center was set as the spot boundary. The length deviation between the spots in different series are all less than 0.3 μm, indicating the excellent dimensional accuracy and consistency of Matrix-seq. It should be explained that there was a deviation between the average size and the design size, which we suspected was introduced by microchannels and belonged to the normal deviation in lithography processing. The normalized fluorescence intensities of spots indicated that the intensity uniformity between spots was better than 3.1%, which was crucial to truly reflect the expression levels of different spatial regions. It is worth emphasizing that there is no coffee ring effect, which deteriorates the uniformity of reactants distribution in droplets and often need to be suppressed by adjusting the composition of reactants and conditions, making this method more friendly to the production environment and operation. To analyze the relationship between spot size and the number of cells, the DAPI-stained MOB section was merged with the three matrices (fig. S2). When the spot size is as low as 10 μm, the spatial resolution can reach the single-cell level (Fig. 2F).

The sequencing data were processed by umitools with a custom protocol to extract the UMI, BC-X and BC-Y in each pair of reads (*18*). The final spatial barcodes (BC − *X*_i_*Y*_j_) were concatenated from the two. The processed reads were trimmed and aligned against the mouse gene reference by STAR (*19*). The featureCounts (version 2.0.3) was used to count the UMIs of each gene for evaluation of capture performance (*20*). Due to the different spot size between different technologies, in order to evaluate the capture ability fairly, we chose the number of gene/UMI per unit area (*μm*^2^) as the standard (Fig. 2G). Except for DBiT-seq using a fixed embryonic brain, other methods used fresh frozen MOB as the tissue sample. Matrix-seq exhibits strong capture performance on the same level as Visium (Fig. 2G), indicating that the two-stage reaction do not deteriorate the capture performance. The number of UMIs captured by this method is lower than that of the others, which is due to the unsaturation of the sequencing depth.

### Spatial transcriptomic mapping of MOB

To test the performance of Matrix-seq, we first profiled the MOB, which is the model tissue widely used in ST approaches(*6, 14, 21*). A matrix with 4900 barcoded spots of 50 μm was used and capture numbers ranging on average from 2611 genes and 5085 transcripts per spot (fig. S4). The H&E image and mapping of UMI of the same tissue section were shown in Fig. 3A. Compared to separate analysis on multiple adjacent sections, this method performing staining-based microscopic imaging analysis and ST analysis on the same section provides the most accurate data for joint analysis, which is very important for understanding the pathology and function of a specific area. We selected two genes (Pcp4 and Slc17a7) as significantly expressed genes to assess the correlation between expression levels and specific structures. The distribution of the two genes shown in Fig. 3B and 3C were remarkable similarity to the ISH results from Allen Brain Atlas(*22*). It is worth mentioning that although the matrix was modified by two-stage reactions, due to the excellent uniformity, the irregular boundaries of structures in the tissue were truly reflected, and the expression level in adjacent spots on same structures maintains a good continuity. To assess the potential diffusion problems during permeabilization, we traced the fluorescent cDNAs to show a pattern of captured transcripts (fig. S5). The fluorescent cDNAs were strictly localized directly under cells. It can be seen in the detailed images that although there was a diffusion effect at the micron level, which did not affect the resolution of 50-μm matrix, but it is meaningful to guide the subsequent development of high-precision matrix. Then, unsupervised clustering was performed to computationally reconstruct the spatial identity (Fig. 3D and 3E). Based on the H&E image and general histology, the clusters were annotated as the corresponding anatomical structures, including the olfactory nerve layer (ONL), subependymal zone (SEZ), granular cell zone deep (GCL-D), granular cell layer internal (GCL-I), granular cell layer externa (GCL-E), internal plexiform layer (IPL), mitral layer (ML), outer plexiform layer (OPL), glomerular layer (GL), accessory olfactory bulb (AOB) and damaged area (DA). Tissue structures with only a single spot width, such as ML, can be resolved, which also proves the excellent resolution of this technique. Further, analysis of differentially expressed genes (DEG) was used to show the gene expression heatmap of clusters (Fig. 3F).

**Fig. 3.**
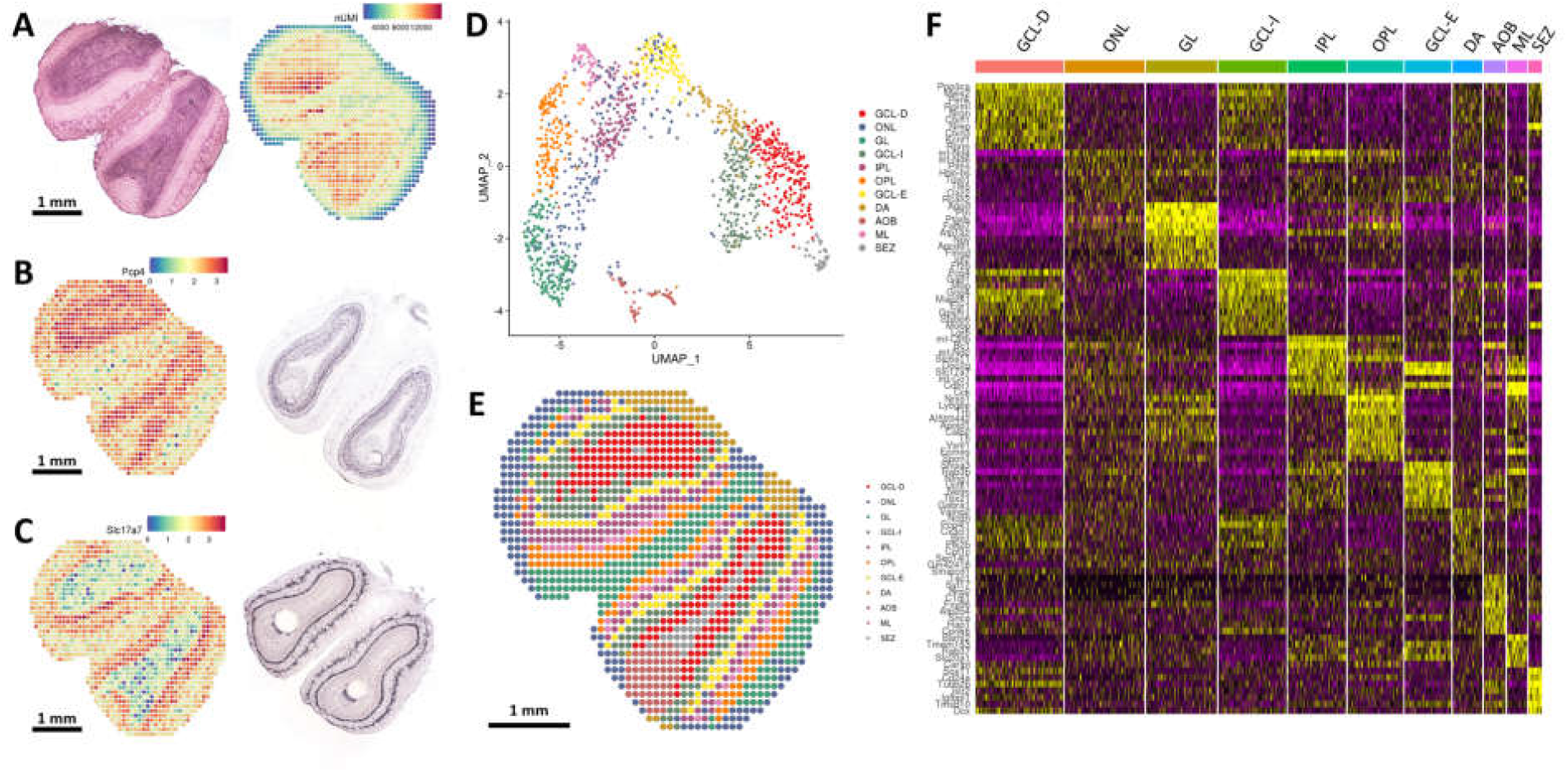
Spatial transcript mapping of MOB. **(A)** H&E image of the MOB section and heatmap of UMI number captured from the same section. **(B)(C)** Spatial heatmap of significant gene expression (Pcp4 and Slc17a7). The ISH results are showed in left. In the high expression region, the two showed excellent consistency. **(D)(E)** UMAP analysis and spatial distribution of 11 distinct clusters classified by unsupervised clustering, which are annotated according to the histological structures. **(F)** Gene expression heatmap of 11 clusters. Top ranked differentially expressed genes are shown in each cluster.

### Spatial transcriptomic mapping of mouse brain

To further test the performance of Matrix-seq in complex tissues, we used the matrix slide with 50-μm spots to analyze a half mouse brains section. Characterizing the gene expression and spatial position of diverse type cell in the brain is fundamental to understand normal brain function and the mechanisms of neural diseases, such as neurodegenerative disorders. The ST technology promises new insights into the relationship between genes, brain and behavior(*22*). The matrix captured 3459 genes and 7690 UMIs per spot on average (fig. S6). The H&E image and mapping of UMI of the same tissue section were shown in Fig. 3A and 3B. Unsupervised clustering also was used here and identified 11 spatial clusters. It is worth emphasizing that the pyramidal layer (cluster-10) was identified, which was a thin layer with a thickness of only tens of micrometers and appeared as a filament with a single spot width in the section. In addition, the multi-layered structure of the neocortex was also well identified. We further visualized selected marker genes to verify the consistency between the spatial distribution of gene expression detected by Matrix-seq and histological structure. The genes Mef2c, Pcp4, Mobp and Hpca are known makers for the neocortex, thalamus, fiber tracts and hippocampus, respectively(*23*). The results in Fig. 4D demonstrated this technique can truly detect the gene expression of different cell types in tissue sections with high spatial resolution and sensitivity.

**Fig. 4.**
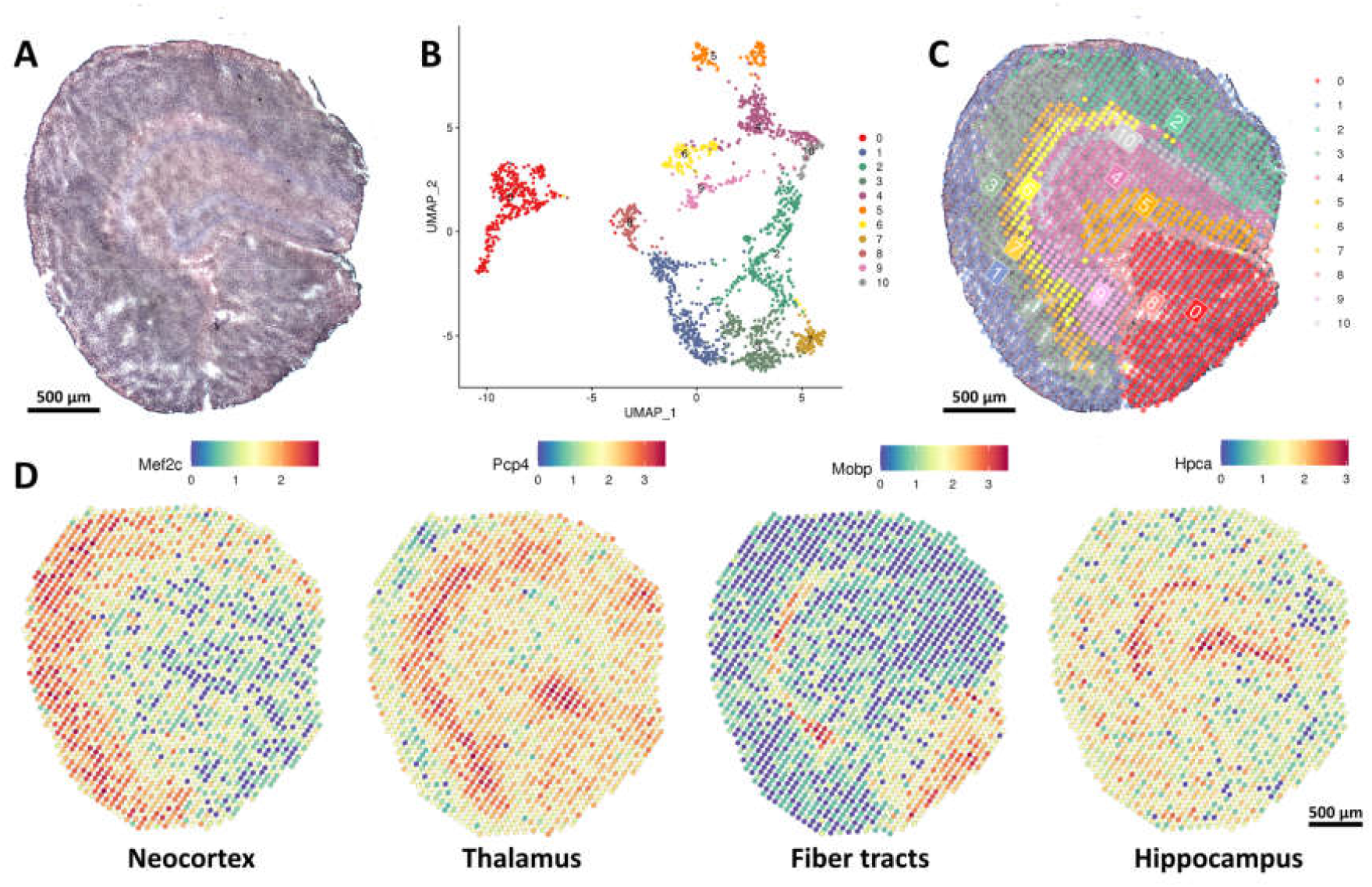
Spatial transcript mapping of mouse brain. **(A)** H&E image of the half mouse brain section. **(B)** UMAP analysis of spots. They are classified into 11 clusters. **(C)** Spatial distribution of the distinct clusters annotated by unsupervised clustering. **(D)** Spatial heatmaps of marker genes. The spatial heatmaps show the position and expression of selected genes (Mef2c, Pcp4, Mobp and Hpca) in the section, which is consistent with the histological structures based on the H&E image.

## Discussion

Matrix-seq innovatively introduces the idea of matrix coding in the preparation of the ST slides, and realizes the modification of barcoding primers on the substrate through the two-stage reaction based on the two types of microfluidics. High-coverage and high-precision matrices were fabricated to verify the potential of the technology. To further facilitate adoption of the technology, we generated a barcoded matrix with 4900 spots for RNA capture from the mouse MOB and brain. It is worth emphasizing that this work provides only one paradigm, the Matrix-seq has many more possibilities in terms of processing options and performance. The mature systems of DNA probe immobilization provide high-performance solutions for the first-stage reaction and are compatible with a variety of solid-phase substrates, not only glass slides. In terms of performance, the current states are far from the limit of this technology. By optimizing the number and morphologies of microchannels, there is still room for further improvement in the effective area, coverage and resolution. In addition, conventional soft-lithography technology for four-inch wafer allows the number and length of microchannels to be expanded to achieve a field of view of several centimeters. The combination of microfluidic matrix for spatial indexing, simple operational procedures in line with clinical practice, and easy-to-use bioinformatics analysis process should support wide application of Matrix-seq in both biological research and clinical pathology.

## Materials and Methods

### Regent and resource

| name | source | Cat No. |
| --- | --- | --- |
| Nexterion® Slide E | SCHOTT NEXTERION® | 1066643 |
| T4 DNA Ligase | New England Biolabs | M0202L |
| Methanol, for HPLC, ≥99.9% | Millipore Sigma | 34860 |
| 2-Propanol (Isopropanol), ≥99.5% | Millipore Sigma | I9516-25ML |
| Hematoxylin, Mayer's (Lillie's Modification) | Agilent | S30930-2 |
| Bluing Buffer, Dako | Agilent | CS70230-2 |
| Eosin, Dako | Agilent | CS70130-2 |
| Pepsin | Sigma-Aldrich | P7000-25G |
| Superscript III | Thermo Fisher | 18080085 |
| RNase Inhibitor | Thermo Fisher | AM2694 |
| BSA | NEB | B9000S |
| DMSO | Sigma-Aldrich | 472301 |
| dATP (100 mM) | Thermo Fisher | R0141 |
| dCTP (100 mM) | Thermo Fisher | R0151 |
| dGTP (100 mM) | Thermo Fisher | R0161 |
| dTTP (100 mM) | Thermo Fisher | R0171 |
| Cyanine 3-dCTP | APExBio | B8159-10ul |
| Proteinase K | Qiagen | 19131 |
| PKD buffer | Qiagen | 1034963 |
| Klenow Fragment (3'→5' exo-) | NEB | M0212L |
| KAPA HiFi HotStart ReadyMix Kit | Kapa Biosystems | KK2601 |
| Ampure XP beads | Beckman Coulter | A63880 |
| VAHTS® Universal Plus DNA Library Prep Kit for Illumina V2 | Vazeme | ND627 |

### Oligonucleotides

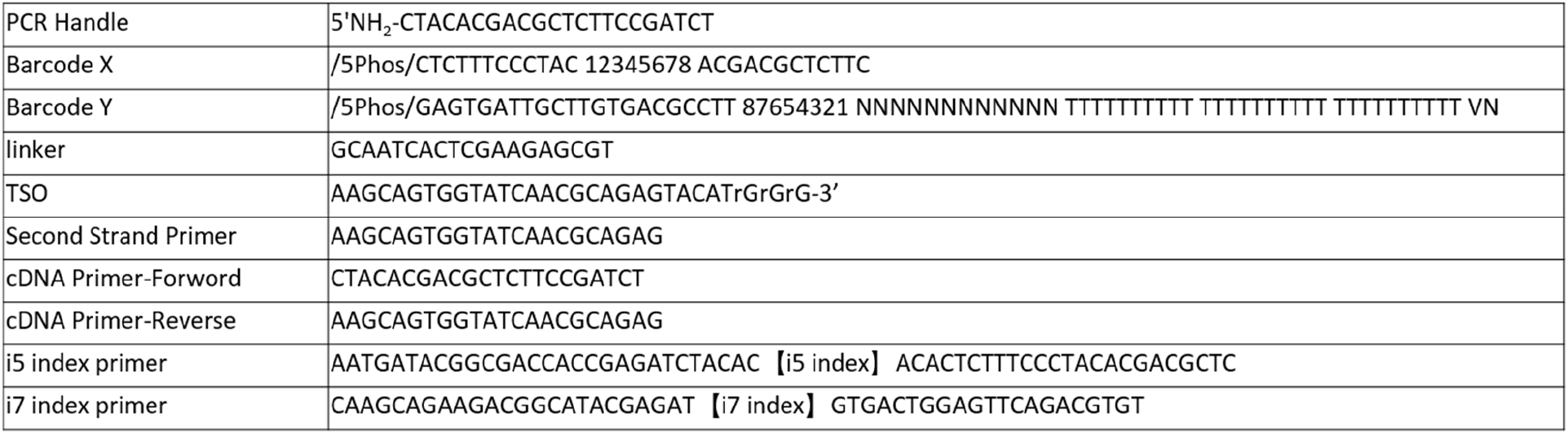

### Fabrication of microfluidic channel

The microfluidic channel was fabricated by standard soft lithography based on Polydimethylsiloxane (PDMS). A thin layer of SU-8 photoresist (Microchem, USA) was spin-coated and patterned by ultraviolet exposure on a silicon wafer. The masks with 10 μm, 25 μm, 50 μm microchannel were ordered from CETC (Nanjing, China). The PDMS base and curing agent (Dow Corning, USA) were mixed in a 10:1 ratio, poured into the SU-8 mold and cured at 65 °C for 45 min. Then, the microchannel was peeled off the mold and cut into the desired shape. A metal punch was used to drill holes in the channel to form the inlets and outlets. Finally, the PDMS microchannel was integrated with the glass slide with a customized holder.

### Generation of spatially barcoded arrays

Barcode X: The arrays of Barcode X were generated onto the surface of Nexterion Slide E. Briefly, microchannel X was attached to the slide and they were clamped by a holder. Add 5 μL oligonucleotides X solution (10∼20 μM in PBS) to inlets (one unique barcoded oligonucleotide for each inlet). The samples entered the microchannel through negative pressure in outlets. The oligonucleotides were immobilized in features of microchannel according to the protocols supported by the manufacturer. Briefly, placed the slides with microchannel in a wet box with saturated sodium chloride at 35°C overnight. Then, the microchannel was removed, and the modified slide was washed sequentially with 0.1 % Triton X-100, 1 mM HCl, 100 mM KCl, and then blocked with 0.1 M Tris (pH 9.0), 50 mM ethanolamine, 0.1% SDS at 50 °C. Finally, the modified slide was blown dry by nitrogen.

Barcode Y: The arrays of Barcode Y were generated on the slide by the ligation reaction. The microchannel Y was attached to the modified slide. The sample (Barcode Y (10∼20 μM in PBS), linker and T4 ligase) was injected into microchannel by negative pressure. Same as before, one unique barcoded oligonucleotide for each inlet. After injection, the assembled slide was incubated at 37 °C for 30 minutes for the ligation reaction. Then, the microchannel was washed with 1×PBS buffer and DI water in sequence and blown dry by nitrogen.

Storage of the modified slides: After the modification, the slides were packaged by the vacuum bag and stored in a 4 °C.

### Tissue preparation

Mice (7 weeks for main olfactory bulbs and 2 weeks for mouse brain) were euthanized. The olfactory bulbs and mouse brains were immediately isolated and washed by precooled PBS buffer. After drying by absorbent paper, the tissues were embedded into cold OCT before sectioning. The tissues were sectioned on a cryostat (−20 °C) at a thickness of 10 μm. The tissue sections were mounted onto the barcoded slide.

### H&E staining and imaging

After incubation at 37 °C for 1 min, the tissue-attached slide was fully immersed in precooled methanol and fixed at -20 °C for 30 min. After drying the slide, 500 μl of isopropanol was added and incubated for 1 min at room temperature. Then, the isopropanol was removed and the slide stand at room temperature for drying. We added 1 ml of hematoxylin, bluing buffer and Eosin in turn to evenly cover the tissue sections, and incubated the slide at room temperature for 7 min, 2 min and 1 min, respectively. The slide was washed by RNase-free Water before changing reagents. Finally, after incubating the slide for 5 min at 37 °C, brightfield imaging was performed.

### Permeabilization and RT

The chamber was assembled on the barcoded slide by a clamp holder. Then, 70 μL permeabilase (0.1% pepsin diluted in 0.1N HCl) was added to the chamber. After incubating at 37°C for 5∼15 min, the permeabilase was removed and the chamber was washed with 0.1×SSC.

Then, 70 μl of RT sample was added, which included: 1x first-strand buffer, 5 mM DTT, 500 μM dNTP, 0.19 μg/μl BSA, 1% DMSO, 20 U/μl Superscript III, 2 U/μl RNase inhibitor and 2.5 μM TSO. The chamber was sealed by the membrane and incubated at 50°C overnight. After the reaction, the RT reaction solution was removed and the chamber was washed by RNase-free Water. Then 70 µl of 0.08 M KOH was added to the chamber and incubated for 5 minutes at room temperature. Remove KOH from the chambers and then washed by RNase-free Water.

### Second strand synthesis and cDNA collection

cDNA second-strand synthesis reaction solution was added into the washed chamber, which included: 1x first-strand buffer, 10U Klenow Exo-, 2.5μM Second Strand Primer. After sealing the chamber, it was placed on a temperature control plate to incubate at 37 °C for 1 hour for cDNA double-strand synthesis. After the reaction, the second-strand synthesis reaction solution in the chamber was discarded, and the chamber was washed by RNase-free Water. Then 35 µl of 0.08 M KOH was added to the chamber and incubated for 10 minutes at room temperature. several 1.5ml centrifuge tubes were prepared and 10 µl Tris (1 M, pH 7.0) was pre-added into them. The sample in the chamber was transferred into the tubes for cDNA collection.

### cDNA amplification and purification

The amplification reaction solution was prepared in a new PCR tube on ice. The PCR reaction solution included: 1×Kapa HiFi Hotstart ReadyMix, 0.8 μM cDNA Forward Primer, 0.8 μM cDNA Reverse Primer, 35 μl cDNA template, with a total volume of 100 μl. Per amplification cycles included: 98 °C for 3 min, then cycled at 98 °C for 15 s, 63 °C for 20 s, 72 °C for 1 min, for 15 cycles. After amplification, 0.6× AMpure XP Beads were used to purify the amplified product. Concentration and length distribution of purified product was identified by the Qubit and Agilent Bioanalyzer High Sensitivity chips, respectively. The concentration and the length of fragment should be between 2-20 ng/μl and larger than 1000 bp.

### Library construction and sequencing

The fragmentation reaction solution includes: 5 μl FEA Buffer V2, 10 μl DNA purified in the previous step, 25 μl ddH2O, 10 μl FEA Enzyme Mix V2, and the total volume is 50 μl. The solution was mixed by pipetting in a precooled tube. Then, the tube was placed in the PCR amplifier, which run the following program: 37 °C for 20 min, 65 °C for 30 min, 4 °C Hold.

The adapter ligation reaction solution includes: 25 μl Rapid Ligation Buffer V2, 50 μl DNA fragmented in the previous step, 15 μl DI water, 5 μl Rapid DNA Ligase V2, 5 μl adapter (10pM), and the total volume is 100 μl. The tube was placed in the PCR amplifier, which run the following program: 20 °C for 30 min, 4 °C Hold.

The library amplification reaction solution includes: 25 μl VAHTS HiFi Amplification Mix, 20 μl DNA prepared in the previous step, 2.5 μl i5 index primer (10pM), 2.5 μl i7 index primer (10 pM), the total volume is 50 μl. The solution was mixed by pipetting and placed into the PRC amplifier, which run the following program: 98 °C 20 s, 63 °C 30 s, 72 °C 20 s for one cycle and a process including 13 cycles. After the amplification reaction, 0.9× AMpure XP was used to purify the amplified product. The constructed library was checked by Qubit and Agilent Bioanalyzer High Sensitivity chip for concentration and length distribution, respectively. The concentration should not be lower than 20 ng/μl and the length of fragments should be distributed between 200-600 bp.

Libraries were sequenced by an illumina NovaSeq 6000 in the PE150 mode.

### Data analysis

To obtain transcriptomics information, the Read 1 was processed by umitool (version 1.1.2) to extract the UMI, Barcode-X and Barcode-Y(*18*). The processed read 1 was trimmed and mapped against the mouse gene reference, mm10 (GENCODE Vm23/Ensembl 98) by STAR (version: 2.5.3a)(*19*). The featureCounts (version 2.0.3) was used to assign gene (*20*) and the digital gene expression matrix for down-stream analysis also was generated by umitools. The barcode X and barcode Y correspond to their location coordinates (row X and column Y). The data of spatial transcriptomics were aligned with the image of H&E staining by manual operation.

The data analysis of tissue sections was carried out with Seurat V4.1.1 following standard procedures(*24, 25*). In short, data normalization, transformation, and selection of variable genes were performed using the SCTransform function with default settings. Principal component analysis (PCA) was performed on the top 3,000 variable genes using the RunPCA function, and the first 20 principal components were used for Shared Nearest Neighbor (SNN) graph construction using the FindNeighbors function. Clusters were then identified using the FindClusters function. We used Uniform Manifold Approximation and Projection (UMAP) to visualize data in a reduced two-dimensional space. To identify differentially expressed genes for every cluster, pairwise comparisons of cells in individual clusters against all remaining cells were performed using the FindAllMarkers function (settings: min.pct = 0.1, logfc.threshold = 0.25). Expression heatmap was then generated using top 10 differentially expressed genes in each cluster.

## Data and materials availability

All data are available in the main text or the supplementary materials.

## Supplementary Materials

**fig. S1.**
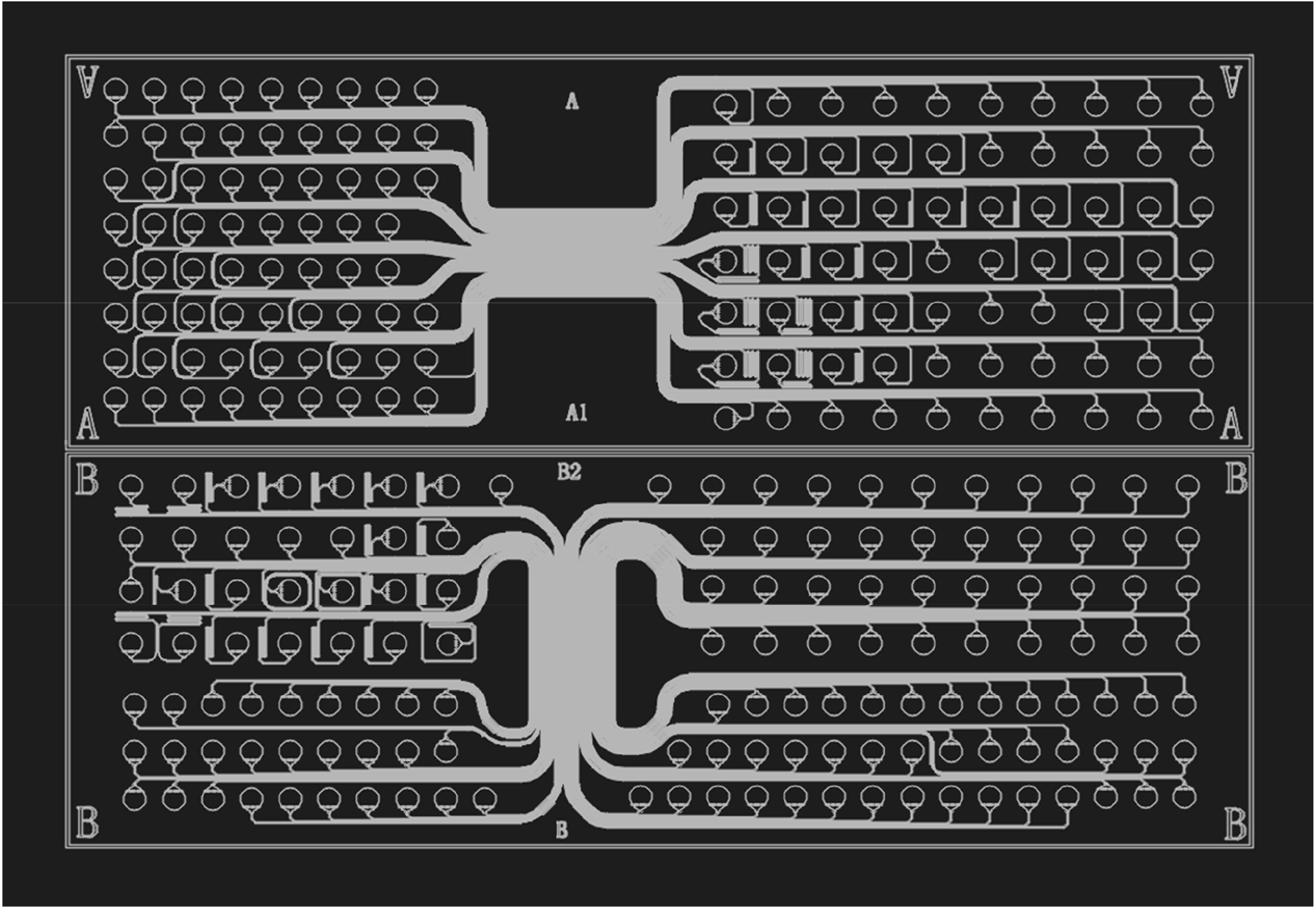
Design diagram of microchannels.

**fig. S2.**
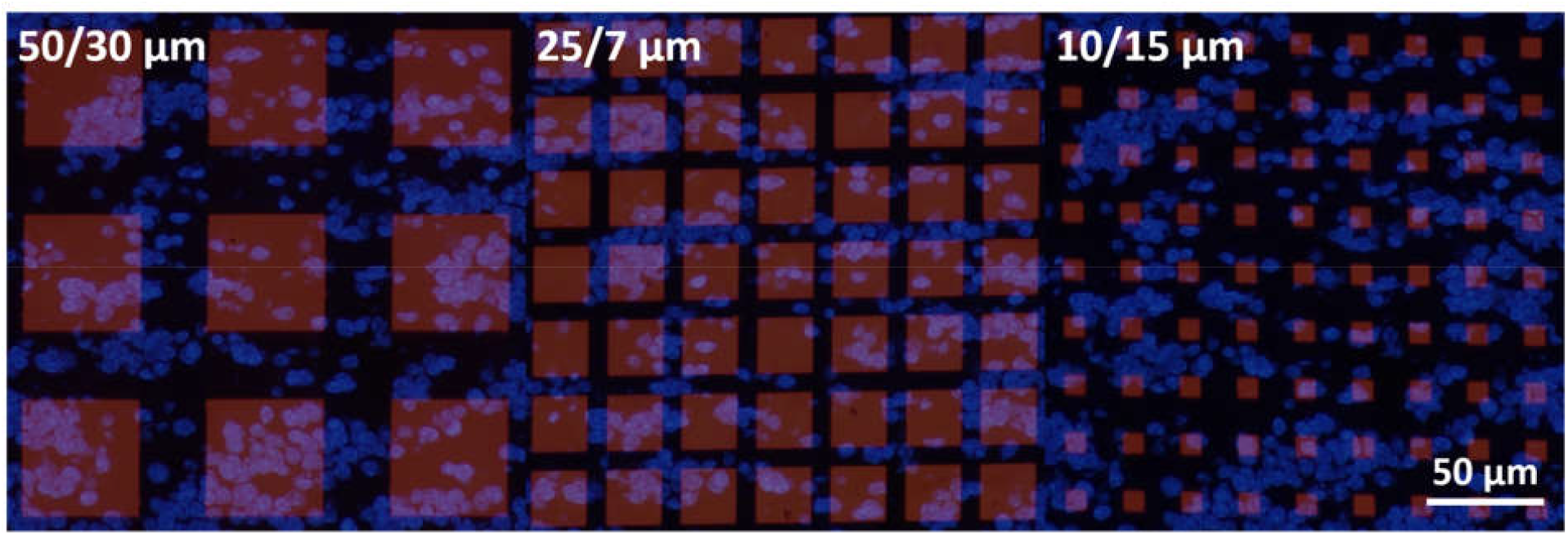
Cell distribution in the matrices with spots of 50, 25, 10 μm. The images are stacked by the images of stained cells and the images of fluorescence-labeled spots. The cells were stained by DAPI and fluorescent spots were labeled by cDNA. cDNA synthesis with Cy3-labeled nucleotides reveals fluorescent cDNA.

**fig. S3.**
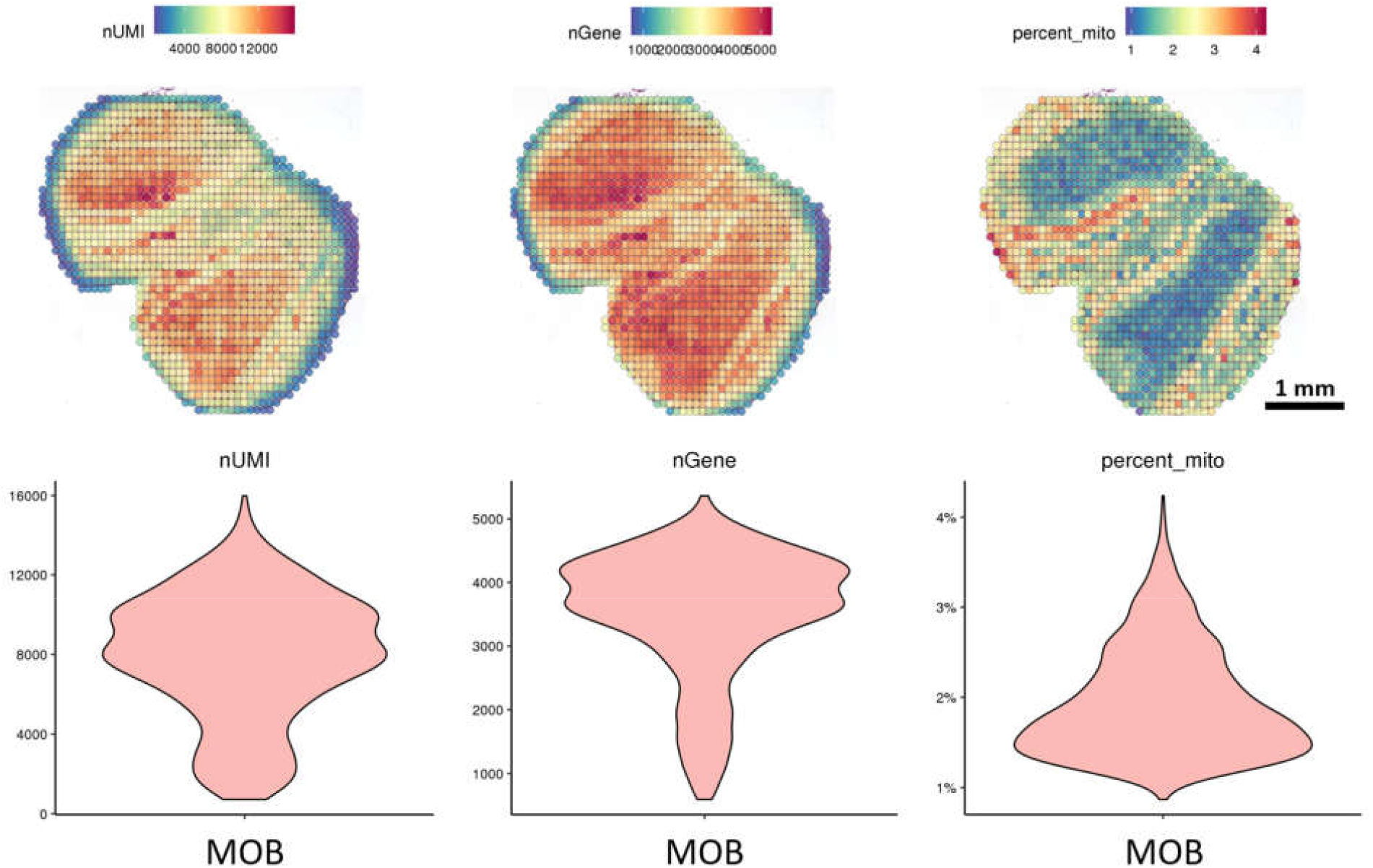
Spatial distributions and statistical graphics of UMI number (nUMI), gene number (nGene) and percentage of mito in MOB.

**fig. S4.**
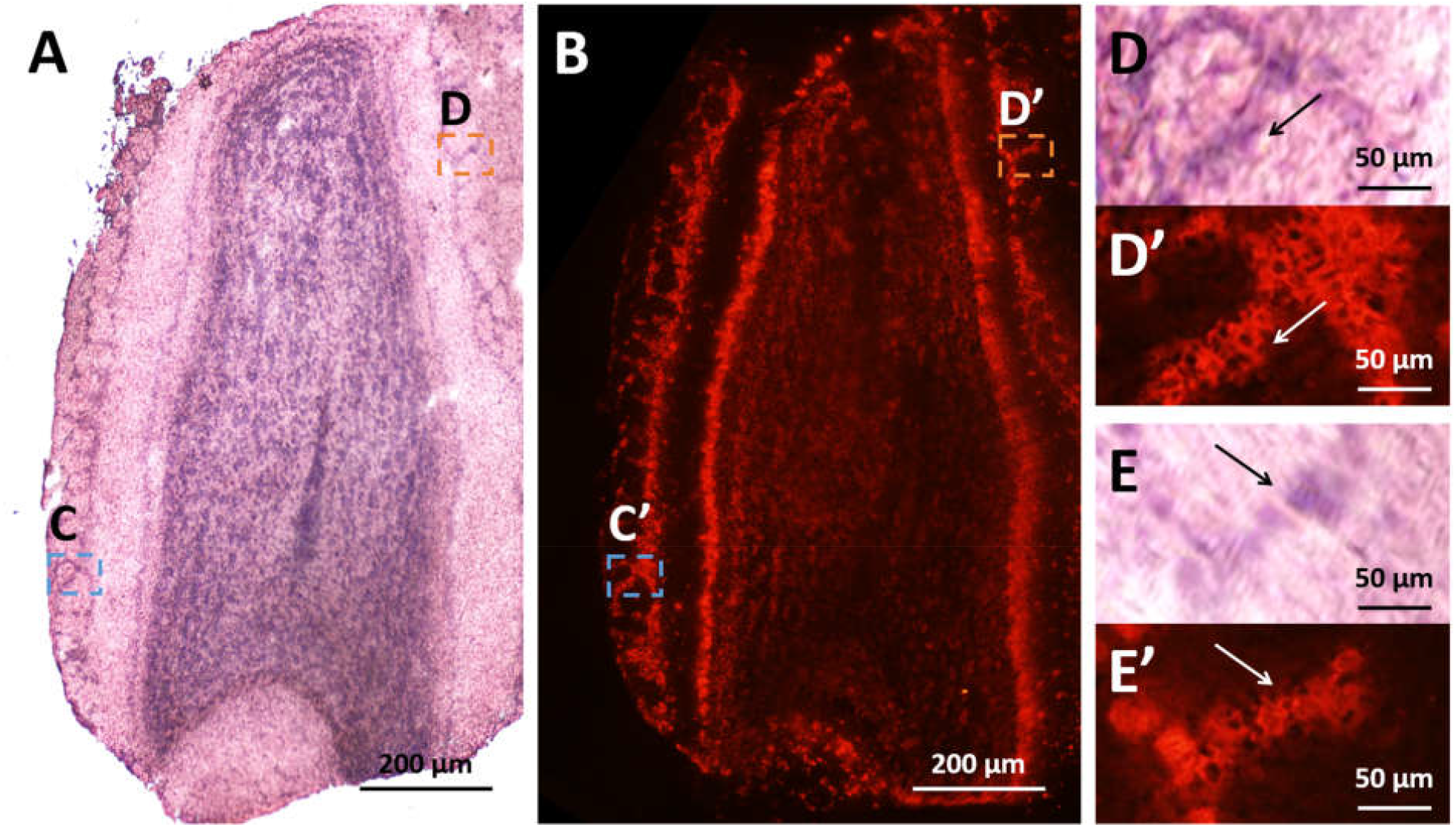
Spatially localized cDNA synthesis. (A) The H&E stain image of the MOB tissue section. (C) Fluorescent image of fluorescence-labeled cDNA after tissue removal. cDNA synthesis with Cy3-labeled nucleotides reveals fluorescent cDNA after tissue removal. (C)(D)/(C′)((D′) Magnified images of cytoplasm/corresponding cDNA.

**fig. S5.**
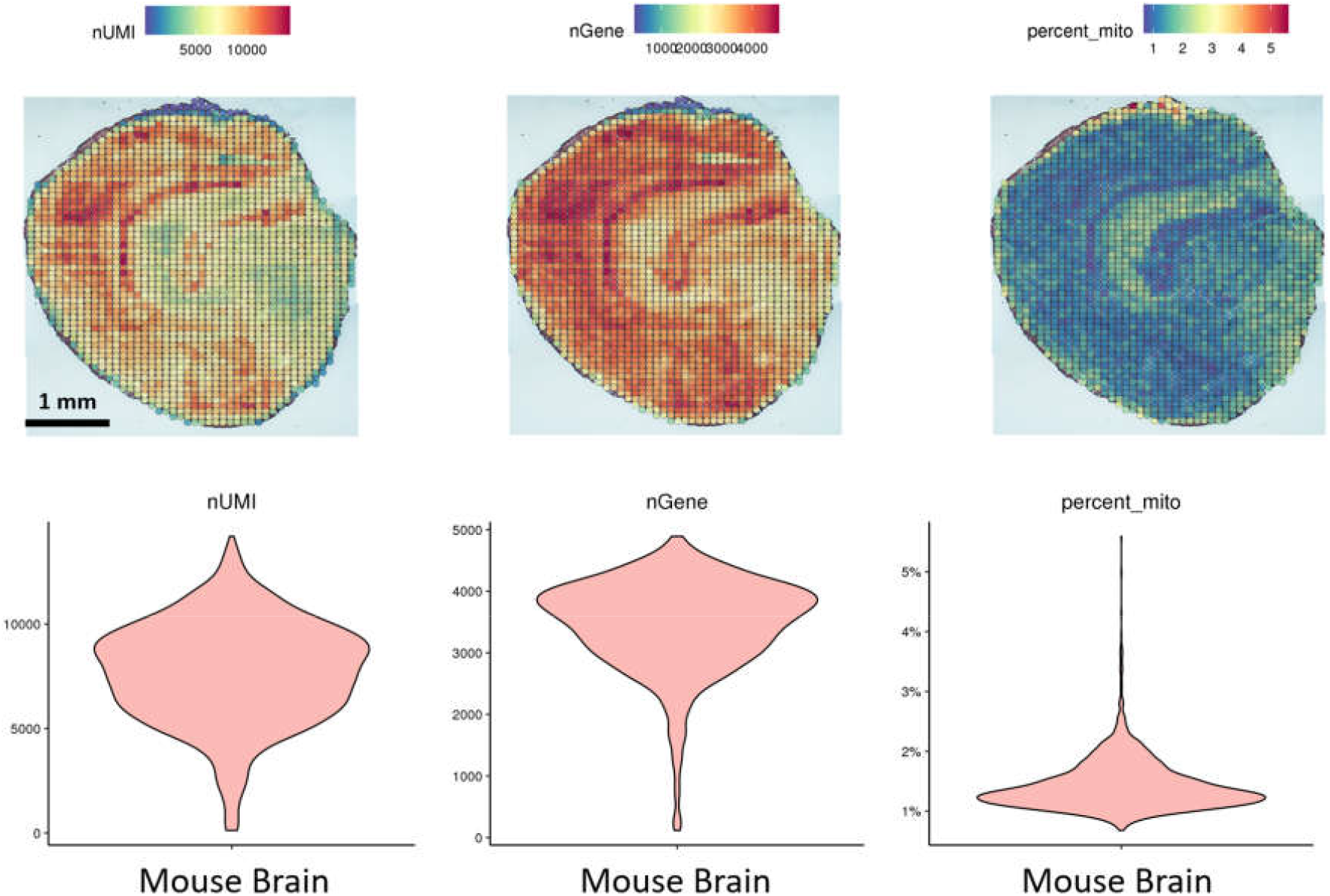
Spatial distributions and statistical graphics of UMI number (nUMI), gene number (nGene) and percentage of mito in the half mouse brain.

